# The small MAF transcription factor MAFG co-opts MITF to promote melanoma progression

**DOI:** 10.1101/2024.09.03.611024

**Authors:** Olga Vera, Michael Martinez, Zulaida Soto-Vargas, Kaizhen Wang, Xiaonan Xu, Sara Ruiz-Buceta, Nicol Mecozzi, Manon Chadourne, Benjamin Posorske, Ariana Angarita, Ilah Bok, Qian Liu, Harini Murikipudi, Yumi Kim, Jane L. Messina, Kenneth Y. Tsai, Michael B. Major, Eric K. Lau, Xiaoqing Yu, Inmaculada Ibanez-de-Caceres, Florian A. Karreth

## Abstract

Transcription factor deregulation potently drives melanoma progression by dynamically and reversibly controlling gene expression programs. We previously identified the small MAF family transcription factor MAFG as a putative driver of melanoma progression, prompting an in-depth evaluation of its role in melanoma. MAFG expression increases with human melanoma stages and ectopic MAFG expression enhances the malignant behavior of human melanoma cells in vitro, xenograft models, and genetic mouse models of spontaneous melanoma. Moreover, MAFG induces a melanoma phenotype switch from a melanocytic state to a more dedifferentiated state. Mechanistically, MAFG interacts with the lineage transcription factor MITF which is required for the pro-tumorigenic effects of MAFG. MAFG and MITF co-occupy numerous genomic sites and MAFG overexpression influences the expression of genes harboring binding sites for the MAFG∼MITF complex. These results establish MAFG as a potent driver of melanomagenesis through dimerization with MITF and uncover an unappreciated mechanism of MITF regulation.

**Significance statement:** MITF is critically involved in melanoma progression and phenotype switching. We discovered that MAFG interacts with MITF to influence expression of MITF target genes and facilitate a shift toward a dedifferentiated melanoma cell state. This study demonstrates that MAFG promotes melanomagenesis by influencing MITF activity, an unappreciated mechanism of MITF regulation.

## INTRODUCTION

Melanoma is an aggressive cancer that affected 331,000 people in 2022, with 60,000 patients succumbing to the disease worldwide^1,2^. The incidence of melanoma has steadily risen over the past 3 decades, with 350,000 new cases and over 70,000 deaths projected for 2025^2^. While resection of early-stage tumors is often curative, metastatic melanoma remains a highly lethal disease. Targeted or immune therapies have improved this dire clinical outlook but are limited in their efficacy by inherent or acquired resistance to these treatments. The plasticity of melanoma cells is critically associated with the development of resistance, enabling phenotype switching in response to environmental cues and stresses^3^. Moreover, melanoma cells can switch between invasive and proliferative states, which fuels the formation of metastasis^3,4^. This phenotype switch has been proposed to be mediated, at least in part, by the microphthalmia-associated transcription factor (MITF), a lineage transcription factor critical for melanocyte differentiation, proliferation, and survival^5,6^. However, the molecular mechanisms driving the phenotype switch in melanoma are incompletely understood.

Activating mutations in BRAF and NRAS occur in >80% of melanomas and significant effort has been invested in exploiting these initiating genetic events and their downstream pathways for therapeutic intervention. Additional genetic events such as loss of the tumor suppressors CDKN2A and PTEN promote the early stages of melanoma progression. However, few mutations specifically driving the advanced stages of melanoma progression and metastasis have been identified^7,8^. Instead, non-genetic mechanisms that influence gene expression programs have emerged as potent drivers of phenotype switching and thus melanoma progression and resistance^9,10^. Transcription factor deregulation enables dynamic and reversible phenotypic adaption, implicating it as a key mechanism of melanoma progression and resistance.

We recently identified the transcription factor MAFG as a bona fide target of the miR-29 melanoma suppressor^11^. MAFG belongs to the small MAF family (sMAFs) of bZIP transcription factors that either homodimerize or heterodimerize with bZIP transcription factors of the Cap’N’Collar (CNC) or BACH families to activate or repress gene expression^12^. Despite their functions as obligate dimerization partners of transcription factors implicated in cancer, most notably the redox and metabolism master regulator NRF2^13^, little is known about the role of sMAFs in human malignancies. In melanoma, MAFG is stabilized by hyperactivation of the MAPK pathway through direct phosphorylation by ERK, facilitating the formation of an epigenetic silencing complex^14^. However, whether increased expression of MAFG promotes the formation of melanoma is unknown.

Here we demonstrate a potent oncogenic effect for MAFG that is associated with a phenotypic switch to less differentiated cell states. MAFG elicits this effect at least in part by binding to and co-opting the lineage transcription factor MITF, a putative driver of melanoma phenotype switching.

## RESULTS

### MAFG overexpression elicits oncogenic effects in vitro and in vivo

We previously reported that MAFG is overexpressed in melanoma cells compared to melanocytes and that its silencing reduces melanoma cell proliferation^11^, prompting us to determine the oncogenic potential of MAFG in melanoma. *MAFG* is amplified, gained and/or overexpressed in 32% of samples in The Cancer Genome Atlas skin cutaneous melanoma (TCGA-SKCM) dataset (**Supplementary Fig. S1A**). Similarly, analysis of two RNA sequencing datasets (GSE3189 and GSE98394) revealed increased *MAFG* in melanoma compared to nevi (**Fig. 1A**). Immunohistochemistry on a tissue microarray containing nevi, primary melanomas, and metastatic melanomas showed elevated MAFG staining intensity in advanced and metastatic samples (**Fig. 1B**). We also observed worse survival in patients with high MAFG expression in the TCGA cutaneous melanoma dataset (**Fig. 1C**). These findings indicate that MAFG is upregulated during melanoma progression and suggest a potential oncogenic role for MAFG. To test this, we overexpressed MAFG in a human immortalized melanocyte (Hermes1) and four human melanoma cell lines (WM164, SKMel28, A375 and WM793) (**Supplementary Fig. S1B**). MAFG overexpression in Hermes1, WM164 and SKMel28 significantly increased cell proliferation (**Fig. 1D**), focus formation (**Fig. 1E**) and, in the case of WM164 and SKMel28, tumor growth in xenograft assays (**Fig. 1F-1H**). In contrast, MAFG overexpression had no effect on A375 and WM793 cells (**Supplementary Fig. S1C and S1D**), suggesting context-dependent oncogenic properties of MAFG. To further characterize the oncogenic role of MAFG in melanomagenesis, we used our embryonic stem cell-genetically engineered mouse modeling (ESC-GEMM) platform^15^ to generate mice harboring a Cre- and Dox-inducible MAFG overexpression allele on a Braf^V600E^; Pten^FL/WT^; Tyr-CreERt2; CAGs-LSL-rtTA3 (BP^MAFG^) background. 4-OHT administration to induce melanomagenesis and feeding a Dox diet to activate melanocyte-specific MAFG overexpression (**Supplementary Fig. S1E and S1F**) drastically accelerated melanoma development (**Fig. 1I**) and reduced overall survival (**Fig. 1J**) compared with a GFP-expressing (BP^GFP^) control cohort. In addition, BP^MAFG^ mice developed significantly more tumors (**Fig. 1K**) and these tumors on average grew moderately faster (**Fig 1L**). Accordingly, at endpoint, there was a trend toward increased Ki67-positive proliferating cells (**Supplementary Fig. S1G**). Tumors were relatively circumscribed masses composed predominantly of small, spindle-shaped cells with focal pigmentation. Tumors with MAFG overexpression had notably more nodular regions exhibiting increased cellularity. (**Supplementary Fig. S1E**). We observed no overt metastases in BP^MAFG^ or BP^GFP^ mice, indicating that MAFG promotes early progression but cannot induce metastasis in the BP model that lacks inherent metastatic propensity. To study whether the tumor suppressor background influences the effects of MAFG, we generated Braf^V600E^; Cdkn2a^FL/FL^; Tyr-CreERt2; CAGs-LSL-rtTA3 mice harboring the same MAFG allele (BCC^MAFG^). Notably, BCC^MAFG^ mice showed decreased survival and increased tumor numbers compared to BCC^GFP^ mice **(Supplementary Fig. S1H-S1J**). Because MAFG is stabilized by MAPK signaling^14^, we also tested if MAFG overexpression is sufficient to induce melanoma in the absence of mutant BRAF. To this end, we generated Pten^FL/FL^; Tyr-CreERt2; CAGs-LSL-rtTA3 mice harboring the MAFG allele (PP^MAFG^). Neither PP^MAFG^ nor PP^GFP^ control mice developed tumors (**Supplementary Fig. S1K**), demonstrating that MAFG overexpression promotes melanoma progression but is insufficient for melanoma initiation.

**Figure 1:**
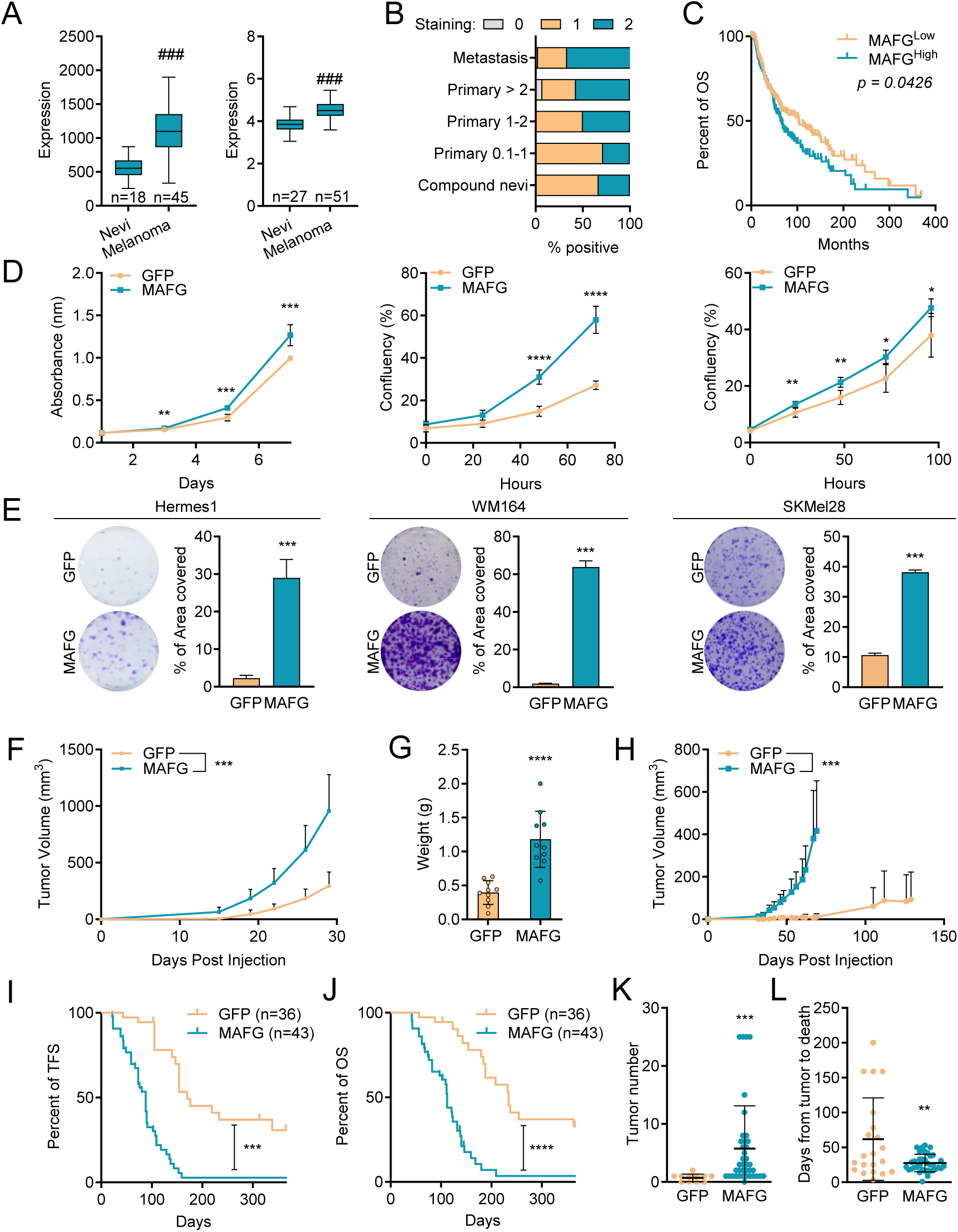
MAFG overexpression elicits oncogenic effects in melanocytes, melanoma cells and in in vivo models of melanoma. (**A**) Expression of *MAFG* in 18 nevi and 45 melanomas obtained from the GSE3189 dataset (left) and 27 nevi and 51 melanomas obtained from the GSE98394 dataset (right). (**B**) MAFG immunohistochemistry on a tissue microarray containing nevi, primary melanomas, and metastatic melanomas where staining of MAFG was graded from 0 (no staining) to 2 (high staining). (**C**) Survival analysis from TCGA (PanCancer Atlas, n = 473) comparing melanoma patients with High or Low expression of MAFG according to the average of normalized read count (cut off point: 1,385.17) using the Log-rank (Mantel-Cox) test. (**D,E**) Proliferation (D) and colony formation (E) assays of Hermes1 (left), WM164 (center) and SKMel28 (right) cells constitutively overexpressing MAFG or a GFP control. (**F,G**) Tumor volume (F) and tumor weight at endpoint (G) of a xenograft assay performed with WM164 cells overexpressing MAFG or a GFP control. (**H**) Tumor volume of a xenograft assay performed with SKMel28 cells overexpressing MAFG or a GFP control. (**I,J**) Kaplan-Meier curves comparing the tumor free survival (I) and overall survival (J) of BP^GFP^ (n = 36) or BP^MAFG^ (n = 43) experimental chimeras using the Gehan-Breslow-Wilcoxon test. (**K**) Number of melanomas that developed in the experimental chimeras of the different genotypes. (**L**) Days elapsed from tumor emergence to when the mice had to be euthanized due to the size of the primary melanomas. For in vitro experiments, representative replicates of at least two independent experiments performed in quadruplicate are shown. * p < 0.05; ** p < 0.01; *** p < 0.001; **** p < 0.0001; ### false discovery rate (FDR) < 0.001.

### MAFG drives a phenotype switch

We next sought to identify the mechanism by which MAFG promotes melanoma. To this end, we performed RNA sequencing analysis of WM164, SKMel28, and A375 cells expressing either MAFG or GFP. Interestingly, while MAFG induced significant transcriptional changes in WM164 and SKMel28 cells (926 and 598 differentially expressed genes, respectively), it had a modest impact on gene expression in A375 cells (167 differentially expressed genes) (**Supplementary Fig. S2A**), corroborating the results of the cell biological assays (**Supplementary Fig. S1C and S1D**). Despite MAFG being an obligate binding partner of NRF2, MAFG-induced differentially expressed genes in WM164 cells did not significantly overlap with previously reported NRF2 expression signatures (**Supplementary Fig. S2B**). NRF2 silencing in WM164 cells overexpressing MAFG (**Supplementary Fig. S2C**) did not diminish proliferation or colony formation (**Supplementary Fig. S2D and S2E**). Moreover, MAFG overexpression did not increase the expression of a NRF2 transcriptional reporter (6xARE-Luciferase) (**Supplementary Fig. S2F**), indicating that the oncogenic effects of MAFG in melanoma are independent of NRF2.

Analyses of the RNAseq results using various pathway and gene ontology tools identified several cancer-associated pathways. Notably, downregulation of the melanin biosynthesis pathway was detected by several tools (**Fig. 2A and Supplementary Fig. S3A**). Melanin biosynthesis is chiefly regulated by the lineage transcription factor MITF^6,16^. Additionally, MITF plays a role in melanoma phenotype switching where it is expressed in melanocytic states while the receptor tyrosine kinase AXL is expressed in less differentiated cell states^5,17,18^. MAFG overexpression moderately diminished MITF levels (**Fig. 2B and Supplementary Fig. S3B**) while the expression of AXL was induced (**Fig. 2B and 2C**). Moreover, MAFG overexpression reduced the expression of the MITF targets *MLANA*, *TYR*, and *PMEL* (**Fig. 2D**), suggesting a role for MAFG in regulating MITF activity and melanoma phenotype switching. Using previously established gene signatures associated with seven melanoma cell phenotypic states^19^, we observed that MAFG altered the expression of these signature genes. Specifically, the Melanocytic signature was diminished in favor of all other signature (WM164) or the Antigen Presentation and Mesenchymal-like signatures (SKMel28) (**Fig. 2E and 2F**). MAFG expression analysis in human^20^ and mouse^19^ scRNAseq data revealed that fewer cells in the cluster defined by the Melanocytic expression signature expressed MAFG and the expression was lower compared to other clusters (**Fig. 2G and 2H, Supplementary Fig. S3C and S3D**). We also determined the baseline phenotypic states of WM164 and SKMel28 cells as well as of A375 and WM793 cells where MAFG overexpression had no effect. Parental WM164 and SKMel28 cells expressed high levels of MITF and its targets *PMEL* and *TYR* and low levels of AXL while A375 and WM793 cells showed the opposite expression pattern (**Supplementary Fig. S3E and S3F**). Moreover, WM164 and SKMel28 cells showed an enrichment of the signature defining the Melanocytic state while the Mesenchymal-like, Stem-like, and Neural Crest-like states are enriched in A375 and WM793 cells (**Supplementary Fig. S3G**). These results indicate that MAFG upregulation promotes aggressive behaviors of melanoma cells in the Melanocytic cell state by inducing a phenotype switch.

**Figure 2:**
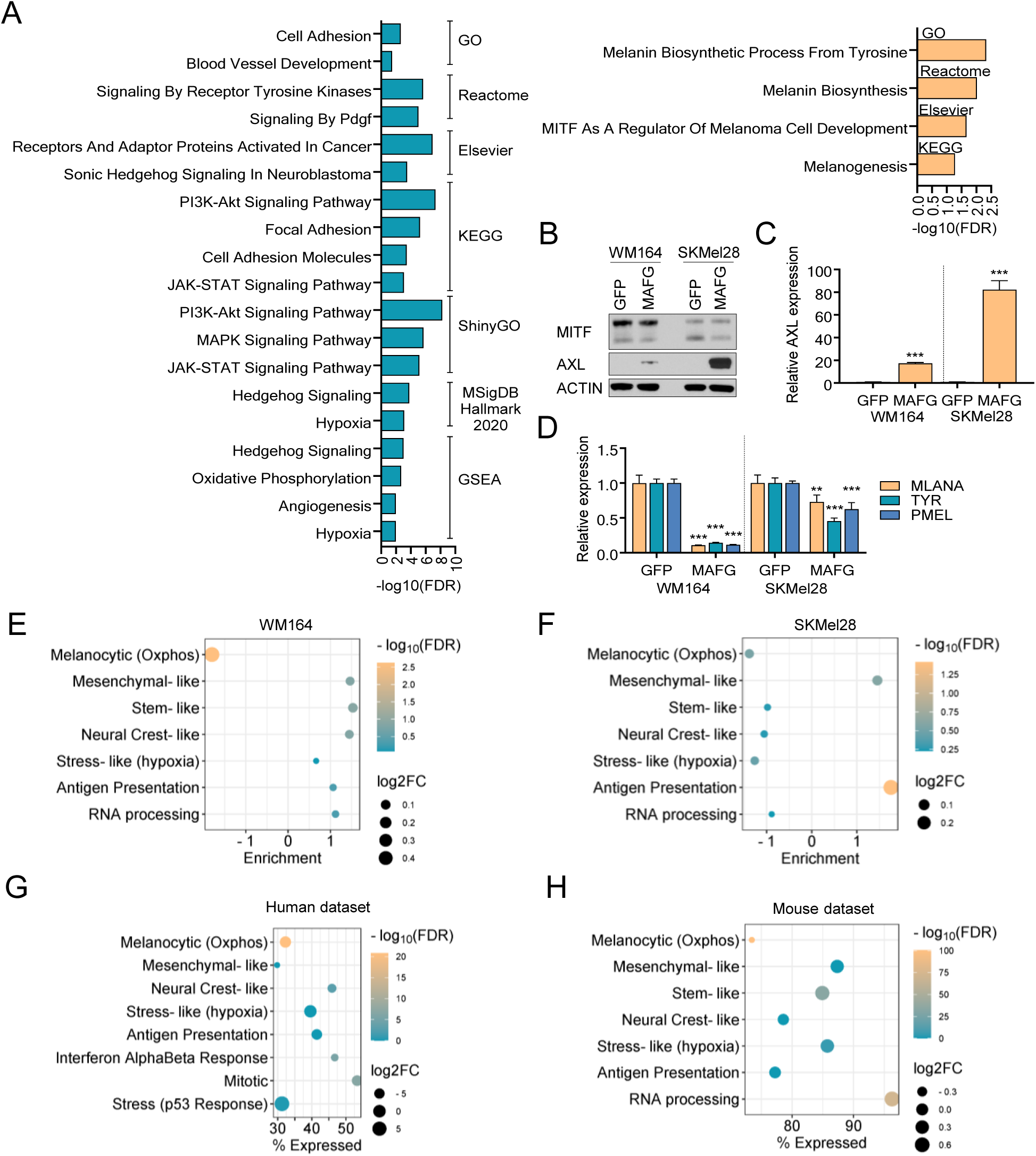
MAFG promotes a phenotype switch. (**A**) Pathway analysis of the upregulated (left) and downregulated (right) genes identified by RNA-sequencing comparing GFP and MAFG overexpression in WM164 cells. (**B**) Expression changes of MITF and AXL by western blot in WM164 and SKmel28 cells overexpressing either GFP or MAFG. (**C**) Expression changes of AXL measured by qRT-PCR in WM164 and SKmel28 cells overexpressing either GFP or MAFG. (**D**) Expression changes of MITF-target genes MLANA, TYR and PMEL measured by qRT-PCR in WM164 and SKmel28 cells overexpressing either GFP or MAFG. (**E,F**) Expression changes of the melanoma phenotype-associated gene signatures in WM164 (E) or SKmel28 (F) cells expressing GFP or MAFG. (**G**) Expression of MAFG in the human melanoma cell clusters defined by cell state gene signatures. (**H**) Expression of MAFG in the murine melanoma cell clusters defined by cell state gene signatures. * p <0.05; *** p < 0.001.

### MAFG interacts with MITF

The modest MITF expression changes induced by MAFG overexpression suggested that MAFG impacts MITF activity through non-transcriptional mechanisms. Interestingly, MITF is a basic helix-loop-helix leucine zipper transcription factor and we hypothesized that MAFG directly interacts with MITF via their leucine zippers. Biotin proximity labeling using an N-terminal TurboID-MAFG fusion construct in WM164 cells (**Supplementary Fig. S4A and S4B**) identified MITF as a putative MAFG interaction partner (**Fig. 3A, Supplementary Table 1**). While MITF did not display the strongest interaction, the fact that this assay identified known MAFG interaction partners such as MAFF, BACH1, BACH2, and NRF1 lent confidence to the notion that MAFG interacts with MITF. We validated the interaction by co-immunoprecipitation of MAFG and MITF in WM164 and SKMel28 cells overexpressing MAFG (**Fig. 3B**). Similarly, we observed the interaction of MAFG with MITF by co-immunoprecipitation in parental WM164 cells expressing endogenous MAFG (**Fig. 3C**). We then performed proximity ligation assays (PLA) in WM164, SKMel28 and A375 cells overexpressing either GFP or MAFG. Nuclear PLA foci were present in WM164 and SKMel28 cells and, to a lesser extent, in A375 cells and the number of foci significantly increased upon MAFG overexpression in all three cell lines (**Fig. 3D and Supplementary Fig. S4C**). We also performed PLAs on a melanoma metastasis TMA. This revealed the presence of nuclear PLA foci in human melanoma specimens (**Fig. 3E**) and showed a correlation between the number of PLA positive nuclei and the number of PLA foci per cell (**Supplementary Fig. S4D**). These results strongly indicate a direct interaction of MAFG with MITF in melanoma.

**Figure 3:**
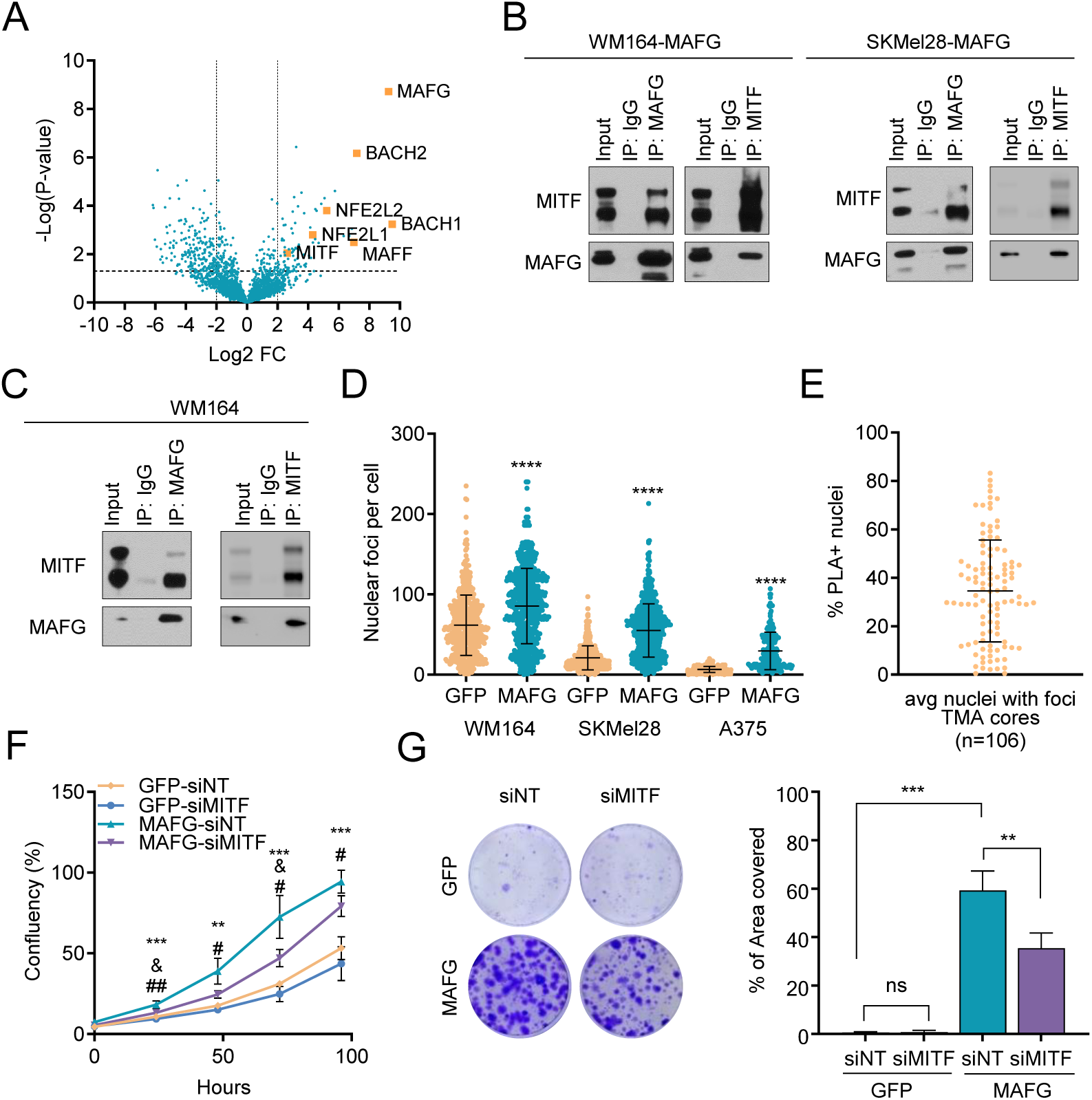
MAFG interacts with MITF. (**A**) Volcano plot showing enrichment of biotin-labeled proteins in WM164 cells expressing N-terminal MAFG-TurboID or control TurboID. Canonical interaction partners of MAFG and MITF are highlighted. (**B**) Co-immunoprecipitation of MAFG and MITF in WM164 and SKMel28 cells overexpressing MAFG. (**C**) Co-immunoprecipitation of endogenous MAFG and MITF in parental WM164 cells. (**D**) Quantification of the proximity ligation assays (PLA) from Fig. S4 in WM164, SKMel28 and A375 cells overexpressing either GFP or MAFG. (**E**) Quantification of PLAs on a melanoma metastasis TMA. (**F,G**) Proliferation (F) and colony formation (G) assays in WM164 cells overexpressing MAFG or GFP transfected with siMITF or control siRNAs. For in vitro experiments, representative replicates of at least two independent experiments performed in quadruplicate are shown. GFP-siNT vs MAFG-siNT: ns, not significant; ** p < 0.01; *** p < 0.001; **** p < 0.0001. GFP-siNT vs GFP-siMITF: & p < 0.05. MAFG-siNT vs MAFG-siMITF: # p<0.05; ## p < 0.01.

To determine whether MITF is required for the oncogenic effects elicited by MAFG, we silenced MITF in WM164 and SKMel28 using a SMART pool of 4 siRNAs (**Supplementary Fig. S5A**). MITF expression was required for the growth of SKMel28 control cells expressing GFP (**Supplementary Fig. S5B and S5C**), precluding conclusions as to whether MITF is required for the effects of MAFG. However, WM164 control cells tolerated the silencing of MITF, and we observed a significant reduction of MAFG-induced proliferation and focus formation (**Fig. 3F and 3G**). Using Dox-inducible shRNAs as an orthogonal approach of MITF silencing in WM164 cells showed similar reductions of MAFG-induced proliferation and focus formation (**Supplementary Fig. S5D-F**). These results indicate that MITF is required for the oncogenic effects of MAFG in melanoma.

### The interaction with MAFG impacts MITF target gene binding and transactivation

To map MAFG and MITF binding sites across the genome, we performed CUT&RUN on WM164 cells overexpressing MAFG or GFP control. Sequencing of DNA fragments identified 24,424 peaks for MAFG and 20,457 peaks for MITF (**Fig. 4A**) that were primarily located in promoters and introns (**Fig. 4B**). We then determined which genes associated with MAFG peaks or MITF peaks are differentially expressed upon MAFG overexpression using the RNA-seq dataset (**Supplementary Fig. S2A**) and performed gene set enrichment analysis. This revealed pathways such as Protein digestion and absorption, Axon guidance, and Rap1 signaling pathway for both MAFG peak-associated genes and MITF peak-associated genes (**Fig. 4C, 4D**), suggesting a certain overlap of genes associated with MAFG and MITF peaks. Indeed, 13,288 of MAFG and MITF peaks overlapped (**Fig. 4A, Supplementary Table 2**), demonstrating that MAFG and MITF bind similar genomic sites.

**Figure 4:**
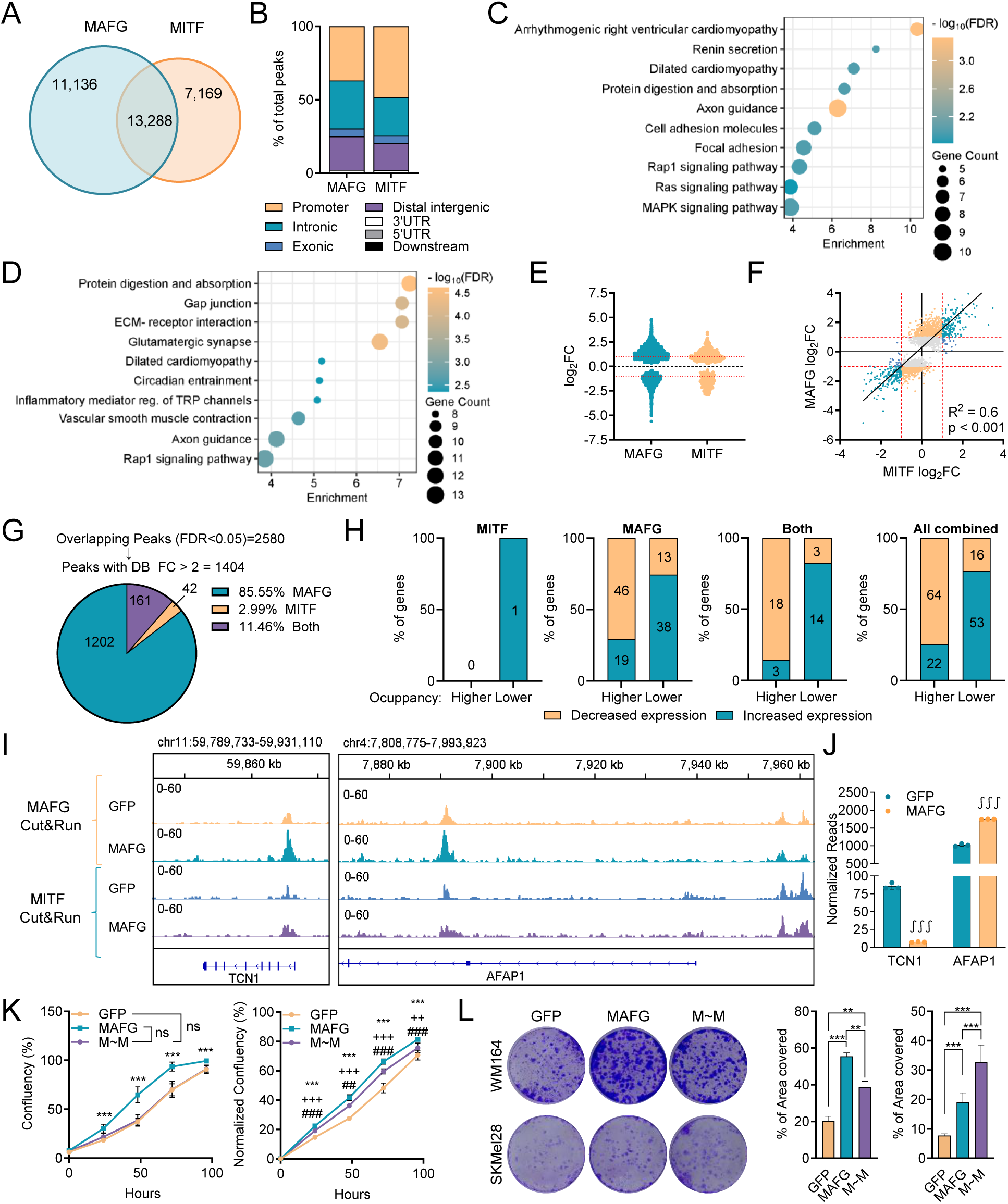
MAFG impacts MITF target gene binding and transactivation. (**A**) Venn diagram showing the overlap between CUT&RUN peaks identified for MAFG and MITF in WM164 cells. (**B**) Distribution of the MAFG and MITF peaks identified in (A) according to their genomic location. (**C,D**). Gene Set Enrichment Analysis (GSEA) of genes harboring MAFG (C) or MITF (D) binding sites and that are differentially expressed in response to MAFG overexpression in WM164 cells. (**E**) Effect of MAFG overexpression on genome binding of MAFG and MITF. The log2FC for MAFG and MITF peaks demonstrating differential binding with FDR < 0.05 is shown. (**F**) Correlation of the differential MAFG and MITF binding at genomic sites having overlapping MAFG and MITF peaks from (E). (**G**) The number of genomic sites having overlapping MAFG and MITF peaks and that exhibit more than 2-fold differential binding for MAFG, MITF or both is shown. (**H**) Percentage of differentially expressed genes with peaks that show differential occupancy of MITF, MAFG, both or all combined peaks from (G). (**I**) IGV plots for *TCN1* (left) and *AFAP1* (right), which harbor overlapping peaks and where MAFG overexpression increases MAFG occupancy without significantly affecting MITF occupancy. (**J**) Expression levels of genes in (G) from the RNAseq results shown in Figure 2. ∫∫∫ false discovery rate (FDR) < 0.001. (**K, L**) Proliferation (K) and colony formation (L) assays in WM164(left) and SKMel28 (right) cells overexpressing a MAFG∼MITF dimer (M∼M) or a GFP control. For in vitro experiments, representative replicate of at least two independent experiments performed in quadruplicate are shown. Ns, not significant; ** p < 0.01; *** p < 0.001. GFP vs M∼M: ++ p <0.01; +++ p <0.001. MAFG vs M∼M: ## p < 0.01; ### p < 0.001.

To determine whether MAFG overexpression influences chromatin binding of MAFG and MITF, we first analyzed the global effects of MAFG overexpression on MAFG and MITF peaks. 5,502 MAFG peaks showed differential occupancy (p_adj_ < 0.05), of which 2,696 and 1,363 showed greater than a 2-fold increase and decrease, respectively (**Fig. 4E, Supplementary Table 3**). In contrast, only 425 peaks showed differential MITF occupancy with 145 and 124 peaks exhibiting greater than a 2-fold increase and decrease, respectively (**Fig. 4E, Supplementary Table 3**), suggesting modest effects on MITF chromatin binding upon MAFG overexpression. We then interrogated how MAFG overexpression impacts MAFG and MITF chromatin binding at the 13,288 shared genomic sites. There were 2,580 sites where MAFG overexpression led to differential occupancy (p_adj_ < 0.05) of MAFG and/or MITF. Of those sites, 2,237 sites showed differential occupancy of only MAFG, 76 sites showed differential occupancy of only MITF, and 267 sites showed differential occupancy of both transcription factors (**Supplementary Table 4**). Plotting the log2FC of differential MAFG and MITF occupancy at each of these sites revealed that changes in binding generally correlate between MAFG and MITF (**Fig. 4F**). However, considering only those genomic sites where MAFG and/or MITF occupancy was affected more than 2-fold identified 1,404 genomic sites, with 1,202 sites showing differential occupancy of only MAFG, 42 sites showing differential occupancy of only MITF, and 161 sites showing differential occupancy of both MAFG and MITF (**Fig. 4G, Supplementary Table 4**). These results indicate that overexpression of MAFG strongly affects the binding of MAFG at genomic sites occupied by MITF, while only modestly influencing the occupancy of MITF at those sites.

The 1,404 genomic sites with differential occupancy of MAFG and/or MITF are associated with 1,238 unique genes (**Supplementary Table 5**), of which 735 were annotated in our RNAseq dataset. Considering only genes with significant differential expression (0.58 < log_2_(FC) < −0.58 and p_adj_ < 0.05) revealed that of the 599 genes with differential occupancy of only MAFG, 116 showed differential expression. Of the 126 genes with differential occupancy of both MAFG and MITF, 38 displayed differential expression, and only 1 out of 10 genes with differential occupancy of only MITF showed differential expression (**Fig. 4H**). Notably, increased MAFG occupancy predominantly resulted in decreased gene expression independently of whether MITF occupancy is also affected at the same sites (**Fig. 4H**). Examples of genes with increased MAFG occupancy that showed decreased and increased expression are *SLC7A8* and *TCN1*, and *AFAP1* and *SVIL*, respectively (**Fig. 4I, 4J and Supplementary Fig. S6A, S6B**). MAFG overexpression also directly influenced the expression of canonical MITF targets such as *MLANA* and *TYR* which were both downregulated (**Fig. 2D**). *MLANA* has two sites where MAFG and MITF binding overlaps, and MAFG overexpression led to increased MAFG binding at those sites and modestly increased MITF binding (**Supplementary Fig. S6C**). In contrast, while TYR has several overlapping MAFG/MITF sites, none of them showed differential occupancy for either MAFG or MITF. Instead, MAFG occupancy is increased at a unique site, thereby likely repressing *TYR* expression (**Supplementary Fig. S6D**). Taken together, these results demonstrate that MAFG overexpression influences its chromatin binding at sites occupied by MITF to deregulate gene expression.

To corroborate the functional importance of the interaction of MAFG with MITF, we generated a MAFG∼MITF tethered dimer construct where the MAFG and MITF proteins are fused by a [RS(GGGS)_4_GGRS] linker. This linker has previously been used to fuse MAFG with NRF2 to identify transcription factor binding sites of this dimer^21^. Expression of the MAFG∼MITF dimer (M∼M) in WM164 and SKMel28 cells (**Supplementary Fig. S7A**) enhanced colony formation (**Fig. 4L**) and *AXL* expression (**Supplementary Fig. S7B**), with a more moderate effect on proliferation (**Fig. 4K**). Moreover, RNA sequencing analysis of WM164 cells expressing the M∼M dimer identified 631 differentially expressed genes, 205 of which were also differentially expressed by MAFG overexpression alone (**Supplementary Fig. 7C**). Gene set enrichment analysis on the M∼M RNA-seq data identified pathways and processes (**Supplementary Fig. 7D**) that partially overlapped with the pathways associated with MAFG/MITF occupied genes (**Fig. 4C and 4D**). Importantly, expression of the M∼M dimer resulted in a phenotype switch, with a reduction of the Melanocytic state in favor of the Mesenchymal-like, Stress-like, and Stem-like states (**Supplementary Fig. 7E**). Thus, the M∼M recapitulates the effects of MAFG overexpression, indicating that the oncogenic potential of MAFG is, in part, elicited by its interaction with MITF.

MAFG has 68% and 78% sequence homology with MAFF and MAFK, respectively, with extensive homology in the centrally located Extended Homology Region, Basic Region, and Leucine Zipper^12^. Moreover, knock-out studies revealed functional overlap between the sMAF family members^22^. We therefore tested whether MAFF and MAFK possess oncogenic potential in melanoma cells. Overexpression of MAFF and MAFK in WM164 cells increased AXL expression and moderately decreased MITF levels, similar to the effects observed with MAFG (**Supplementary Fig. S8A**). MAFF and MAFK enhanced proliferation and colony formation (**Supplementary Fig. S8B and S8C**), albeit to a lesser extent than MAFG. The sMAF expression constructs contain dual Myc-Flag tags, enabling the immunoprecipitation with the same antibody. Flag IP of sMAFs readily pulled down MITF (**Supplementary Fig. S8D**), demonstrating that all three sMAF proteins bind MITF. Normalization of MITF to the respective Myc-tagged sMAFs suggested that MAFG binds MITF more strongly than MAFF and MAFK (**Supplementary Fig. S8D**). These data indicate that all three sMAF proteins have oncogenic effects in melanoma, but MAFG is the most potent family member.

## DISCUSSION

Non-mutational mechanisms are emerging as potent drivers of melanomagenesis. Among these mechanisms, transcription factor deregulation enables dynamic and reversible adaptation of transcriptional programs that govern melanoma progression and metastasis. Here, we demonstrate that overexpression of the sMAF transcription factor MAFG potently promotes melanomagenesis at least in part by interacting with MITF and promoting a phenotype switch to a dedifferentiated cell state.

Expression and silencing studies have associated MAFG with several cancer types^11,23,24^ and with the development of platinum resistance of lung and ovarian cancer^25,26^. However, studies that directly tested the oncogenic effects of elevated MAFG expression, especially in an in vivo setting, are limited. Our in vitro and in vivo studies establish MAFG as a bona fide pro-tumorigenic driver in melanoma. Knock-out studies demonstrated functional redundancy between the sMAF family members^12^, and we found that MAFF and MAFK also possess oncogenic potential. However, MAFF and MAFK appear to be less potent in promoting melanoma cell growth despite their ability to interact with MITF. It is possible that MAFG has unique functions that act in concert with its effect on MITF to promote melanoma. Moreover, MAFG harbors a Serine residue that is lacking in MAFF and MAFK and that is phosphorylated by ERK to increase protein stability^14^. The near-universal hyperactivation of the MAPK pathway in melanoma may thus specifically increase MAFG protein levels, which in turn contributes to melanomagenesis. This notion is supported by our finding that MAFG overexpression requires oncogenic BRAF to enhance melanomagenesis.

The fact that MAFG elicits its pro-tumorigenic effects only in the context of an initiating mutation further indicates that MAFG influences melanoma progression. Phenotype switching is emerging as a key facilitator of melanoma progression^27^ and MITF is critically engaged in governing melanoma cell states^3^. Our finding that MAFG induces a dedifferentiated cell state through its interaction with MITF adds to the growing number of molecular mechanisms involved in phenotype switching. Given the prominent role of MITF in phenotype switching, melanocyte differentiation, and pigmentation, it is no surprise that MITF is tightly regulated. This is achieved through extensive transcriptional regulation to control MITF levels as well posttranslational modifications that affect localization, complex formation, and protein stability^6^. Moreover, MITF interacts with co-factors and scaffold proteins that shape its activity^6^. MITF typically dimerizes with members of the MiT/TFE family (MITF, TFEB, TFE3, and TFEC), and atypical dimerization with other transcription factors has also been described. Specifically, MITF may directly interact with β-Catenin and LEF-1^28,29^, suggesting that MITF may influence the output of WNT pathway activation, which, too, has been implicated in melanoma phenotype switching^30^. However, whether the direct interaction of MITF with β-Catenin or LEF-1 affects melanomagenesis remains to be determined. Our finding that the interaction of MAFG with MITF impacts target gene expression and promotes melanomagenesis substantiates an underappreciated mechanism of MITF regulation whereby the formation of atypical heterodimers influences the transcriptional activity of MITF.

sMAF proteins are obligate dimerization partners of CNC/BACH transcription factors, including NRF2^12^. NRF2 may play a role in melanomagenesis^31,32^ but this remains controversial^33^. In the context of MAFG overexpression, NRF2 is dispensable for the oncogenic effects. In colon cancer and melanoma, MAFG recruits a corepressor complex consisting of BACH1, CHD8, and DNMT3B^14,34^ but whether MAFG overexpression enhances promoter hypermethylation through this complex and whether MITF contributes to the activity is unknown. However, it is probable that the interactions of MAFG with the corepressor complex (through BACH1) and MITF are mediated through the respective leucine zippers and are therefore mutually exclusive. This would suggest that the MAFG∼MITF and corepressor complexes form and function independently. While we observed that the tethered MAFG∼MITF dimer elicits oncogenic effects, future studies are needed to evaluate the relative contribution of MITF and the corepressor complex to the oncogenic effects of MAFG.

## MATERIALS AND METHODS

### Cell culture and treatments

The human immortalized melanocytes cell line Hermes1 was obtained from the Functional Genomics Cell Bank at St George’s, University of London, UK, and cultured in RPMI media supplemented with 10% FBS, 10ng/mL hSCF (R&D, Cat # 255-SC), 200nM TPA (Sigma, Cat # P8139), 200pM Cholera Toxin (Sigma, Cat # C8052), and 10nM Endothelin-1 (Sigma, Cat # E7764) at 37°C in a humidified atmosphere containing 10% CO_2_. A375 and SKMel28 cells were purchased from ATCC; WM164 and WM793 cells were a gift from M. Herlyn from the Wistar Collection of Melanoma cell lines. All melanoma cell lines were cultured in RPMI containing 5% FBS at 37°C in a humidified atmosphere containing 5% CO_2_. Lenti-X HEK293T cells were obtained from Takara and cultured in DMEM containing 10% FBS at 37°C in a humidified atmosphere containing 5% CO_2_. All cell lines were routinely tested for mycoplasma using MycoAlert Plus (Lonza, Cat # LT07-710), and human melanoma cell lines were STR authenticated by Moffitt’s Molecular Genomics Core.

### Plasmids

The CMV promoter and puromycin in pLenti-GFP-puro were replaced with the EF1α promoter and blasticidin, respectively, using standard In-Fusion cloning (Takara Bio, Cat. # 638911) to create pLEGB. The Myc-DDK-tagged ORF clone of MAFG (OriGene, Cat. # RC221486) was cloned into pLEGB to replace GFP and create pLEB-MAFG. ORF clones of MAFF and MAFK (OriGene, Cat. # RC215609L3 and #RC223543) were used to replace MAFG in pLEB-MAFG to create MYC-DDK-tagged cDNA expression constructs. TurboID-pCDNA constructs were obtained from E. Padron (Moffitt Cancer Center and Research Institute). MAFG and TurboID-3xNLS were amplified via PCR from the Myc-DDK-tagged ORF MAFG expression plasmid and pCDNA3-TurboID-3xNLS, respectively, and inserted into lentiviral vector pDEST via In-Fusion Recombination. shRNAs targeting MITF were designed using SplashRNA (http://splashrna.mskcc.org/) and cloned into a Doxycycline-inducible vector (pRRL-Puromycin) using the Q5 Site-Directed Mutagenesis Kit (NEB, Cat. # E0554S) with minor modifications. PCR for site-directed mutagenesis was carried out using 2X Platinum SuperFi II Green PCR Master Mix (Thermo Scientific, Cat. # 12369010), following the manufacturer’s recommended three-step protocol. To generate the MAFG-MITF dimer (M∼M), the cDNA of the melanoma-specific isoform of MITF (M-MITF) was first cloned into pLenti-EF1α-Blasticidin followed by cloning the MAFG cDNA downstream of MITF with a flexible linker of 22 amino acids [RS(GGGS)_4_GGRS] using Takara In-Fusion cloning.

### Cell transfection and lentiviral transduction

For siRNA transfections, 100,000 cells/well were plated in 6-well plates and transfected with 25nM of ON-TARGETplus siRNA pool (Dharmacon, NFE2L2 Cat. # L-003755-00-0005; MITF Cat. # L-008674-00-0005) or Non-Targeting control (Dharmacon, Cat. # D-001810-10-05) using JetPrime (VWR, Cat. # 89129-924) according to the manufacturer’s protocol. 4-6 hours after transfection, cells were trypsinized and replated for cell biological assays. For lentiviral transductions, Lenti-X HEK293T cells were transfected with the lentiviral vector and the Δ8.2 and pMD2-VSV-G helper plasmids at a 9:8:1 ratio. Supernatants were collected 48 hours after transfection and filtered through a 0.45μm filter. Hermes1 and melanoma cells were plated in 10cm dishes and transduced with supernatants in the presence of 8 μg/mL polybrene overnight. Selection was carried out by treating cells with 2.5-10 μg/mL Blasticidin for 5 days or 1 μg/mL Puromycin for 3 days.

### Proliferation and focus formation

For proliferation assays with Hermes1, cells were plated in 96-well plates at a density of 4,500 cells/well and harvested for seven days. Cells were fixed and stained with 0.1% crystal violet (VWR, Cat. # 97061-850) solution in 20% methanol for 20 minutes followed by extraction of crystal violet with 10% acetic acid. Absorbance was measured at 600nm using a plate reader. For melanoma proliferation assays, cells were plated in 96-well plates at a density of 1,000 - 2,500 cells/well in 200μL of complete medium. After 24 hours, the plate was loaded into Cellcyte-X live cell analyzer (ECHO). Images of each well were taken daily for 4–5 days and the cell confluency of each image was quantified. For colony formation assays, cells were plated in 6-well plates at a density of 1,000 - 2,000 cells/well and cultured for 2-3 weeks. Cells were fixed and stained with 0.1% crystal violet solution in 20% methanol for 20 minutes. Colonies or area covered was quantified using ImageJ software.

### RNA isolation and quantitative RT-PCR

Total RNA was isolated using TRI-Reagent (Zymo Research, Cat. # R2050-1-200) according to the manufacturer’s recommendations. For qRT-PCR, 500ng of total RNA were reverse transcribed using PrimeScript RT Master Mix (Takara Bio, Cat. # RR036A), and subsequent SYBR Green-based qPCRs were performed as previously described^11^. Primers for SYBR Green qPCR are listed in **Supplementary Table 6**.

### RNA-sequencing

Total RNA from cells was isolated using miRNeasy Mini Kit (Qiagen, Cat. # 217004), quantitated with the Qubit Fluorometer (ThermoFisher Scientific), and screened for quality on the Agilent TapeStation 4200 (Agilent Technologies). The samples were then processed for RNA-sequencing using the NuGEN Universal RNA-Seq Library Preparation Kit with NuQuant (Tecan Genomics, Cat. # 9156). Briefly, 100 ng of RNA was used to generate cDNA and a strand-specific library following the manufacturer’s protocol. Quality control steps were performed, including TapeStation size assessment and quantification using the Kapa Library Quantification Kit (Roche, Cat. # 07960140001). The final libraries were normalized, denatured, and sequenced on the Illumina NovaSeq 6000 sequencer with the SP-200 cycle reagent kit in order to generate approximately 50 million 100-base read pairs per sample (Illumina).The raw RNA-seq reads were first assessed for quality using FastQC (http://www.bioinformatics.babraham.ac.uk/projects/fastqc/). Quality trimming was performed using cutadapt^35^ to remove reads with adaptor contaminants and low-quality bases. Read pairs with either end too short (<25bp) were discarded from further analysis. Next, trimmed and filtered reads were aligned to the human transcriptome GRCh38 using STAR^36^. Uniquely aligned reads were counted at gene level using featureCounts^37^ and then normalized using DESeq2 package^38^ taking into account RNA composition bias. A negative binomial generalized linear model implemented in DESeq2 were used to determine differentially expressed genes. Genes with Fold Change > 2 and false discovery rate (FDR) controlled p-value ≤ 0.05 were considered differentially expressed and visualized using volcano plot and heatmap. The gene list was used to perform pre-ranked gene set enrichment analysis (GSEA^39^ version 4.0.2) to assess enrichment of hallmarks, curated gene sets, and gene ontology^40^ terms in MSigDB^39,41^. We also collected signatures for 7 phenotypic states of melanoma cells^19^ and assessed their enrichment in MAFG overexpression vs. control samples using pre-ranked GSEA. The resulting normalized enrichment score (NES) and FDR controlled p-values were used to assess transcriptome changes. GEO Accession numbers: GSE274945 (token for access: ifelsmmadngbfkl), GSE274811 (token for access: cjytswcanrmxbkl). RNA-seq data of melanoma cell lines^11^ (GSE148552) were analyzed as described above. Gene expression was quantified as Transcript per Million (TPM) using RSEM^42^. Enrichment of the 7 phenotypic states of melanoma cells^19^ was calculated in each sample by single-sample GSEA analysis using GSVA R package. The z-scored enrichment scores were visualized using heatmap.

### CUT&RUN

At approximately 80% confluency, adherent cells were scraped from the plates, washed with PBS, and counted. Isolation of cell nuclei from a minimum of 250,000 cells, and experimental procedures were performed as described using the CUTANA ChIC/CUT&RUN Kit version 4.0 (EpiCypher, Cat. # SKU: 14-1048). Antibodies used were MAFG (Abcam, Cat. # ab154318) and MITF (Millipore Sigma, Cat. # HPA-003259). Following enrichment using the EpiCypher CUT&RUN kit, the samples were quantitated with the Qubit dsDNA HS Assay Kit (Thermo Fisher Scientific, Cat. # Q32851) and 0.6 to 5 nanograms of enriched DNA was used for library preparation using the Kapa HyperPrep Kit (Roche, Cat. # 07962347001) following EpiCypher’s parameters for indexing PCR and library amplification as described in the CUT&RUN Library Prep Manual. The final libraries were Qubit quantitated and screened on the Agilent TapeStation D1000 ScreenTape (Agilent Technologies) to assess the fragment size distribution. Following final quantification using the Kapa Library Quantification Kit (Roche, Cat. # 07960140001), the libraries were sequenced on the NextSeq 500 using a Mid-output 150-cycle Kit in 2×50 configuration to generate 6-8 million read-pairs per sample (Illumina). *FastQC* were used to examine characteristics of the sequencing libraries. Sequence reads of verified libraries were aligned to the human transcriptome GRCh38 using Bowtie2^43^. BigWig files were generated using *bamCoverage* from deepTools^44^ and visualized in IGV. Model-based Alignment (MACS2)^45^ was utilized for peak calling with --q 0.01 using IgG samples as control. A consensus set of peaks was generated by aggregating peaks of individual samples. The human ENCODE blacklist regions^46^ were removed from consensus peaks. Peaks were annotated using *annotatePeak* function in R package ChIPseeker^47^. Number of reads mapped per consensus peak were calculated for each sample using *bamsToCount* function in Rsubread^48^ R package. Raw counts were further normalized by library size and tested for differential expression using DESeq2^38^. Differential peaks were identified as fold-change > 2 and false discovery rate (FDR) controlled p-value < 0.05. Pathway analysis^49^ was further performed using genes harboring differentially expressed peaks. Bubble plots were created using SRplot tool^50^. GEO Accession number: GSE274810 (token for access: glklsywwxbebzmn).

### Public single-cell RNAseq data analysis

We downloaded processed single-cell RNAseq data of mouse melanoma^19^ from https://marinelab.sites.vib.be/en, and processed single-cell RNAseq data of malignant cells of human melanoma metastatic biopsies^20^ from KU Leuven RDR (https://doi.org/10.48804/GSAXBN). Normalized expression of MAFG was visualized on UMAP generated by the original studies. Differential expression of MAFG across different melanoma states of malignant cells was calculated by Wilcoxon signed-rank test, and visualized by log2FC and percentage of expression.

### Biotin Proximity Labeling

TurboID-MAFG and TurboID-NLS expressing cells were cultured in DMEM containing 5% dialyzed FBS at 37°C. For biotin labeling, cells were grown in three 10-cm dishes per experimental condition for 24 h and were given fresh media the day before addition of biotin. At 80-90% confluency, cells were incubated with fresh media and 50 µM biotin in 5% dialyzed FBS in DMEM for 10 min at 37°C. Biotin labeling was stopped by immediately placing cells on ice and five washes with PBS. Cells were scraped and collected to confirm biotinylation of proximal proteins as described previously (DOI: 10.1038/s41596-020-0399-0).

### Sample preparation for mass spectrometry analysis

Cells were sonicated at 4°C in RIPA buffer (10% glycerol, 50 mM HEPES, 150mM NaCl, 2 mM EDTA, 0.1% SDS, 1% Triton X-100, 0.2% Sodium deoxycholate) containing protease inhibitor (Thermo Scientific, Cat. # 78429), phosphatase inhibitor (Thermo Scientific, Cat. # 78426), and benzonase (Sigma, Cat. # E1014-5KU). Protein concentration was measured using Pierce BCA Protein Assay kit (Thermo Scientific, Cat. # 23225). 4.5 mg of protein lysate was incubated with 30 µL of packed pre-washed Streptavidin Sepharose beads (Cytiva, Cat. # 17511301) overnight on a rotator at 4°C. Beads were washed with wash buffer (WB) 1 (2% SDS) twice, once with WB2 (500 mM NaCl, 0.1% deoxycholate, 1% Triton X-100, 1 mM EDTA, 50 mM HEPES, pH 7.5), once with WB3 (250 mM LiCl, 0.5% Triton X-100, 0.5% deoxycholate, 1 mM EDTA, 50 mM HEPES. pH 8.1), and once with WB4 (150 mM NaCl, 50 mM HEPES, pH 7.4). The streptavidin beads were further washed 3 times in 50 mM ammonium bicarbonate (ABC). Beads were then resuspended in 100 µL of 50 mM ABC containing 1 mg trypsin (Promega, Cat. # V5113) and 0.1 mAu Lys-C (Wako Chemicals, Cat. # 129-02541) and incubated overnight at 37°C with shaking for on-bead-digestion. The following day, 0.5 mg trypsin and 0.05 mAu Lys-C were added to the beads and incubated for 2 hours. Digested peptides in the supernatant were collected into a fresh tube and the beads were washed twice with HPLC-grade water and pooled with the peptides. Pooled peptides were centrifuged at 16,000 g for 10 minutes and filtered using BioPureSPN columns (Nest Group, Cat. # C100500), pre-wetted with 0.1% trifluoroacetic acid, and centrifuged at 3,000 g for 2 minutes. Filtered peptides were acidified to 2% formic acid, dried using a speed vac, and stored at −80°C. Peptides were resuspended in 13 µL of 98 parts buffer A (water + 0.1% formic acid) and 2 parts buffer B (99.9% acetonitrile + 0.1% formic acid) and 5 µL of peptides were injected for mass spectrometric analysis.

### Chromatographic separation and label-free quantification

Tryptic peptides were separated by reverse phase nano-HPLC using an Ultimate 3000 RSLCnano System (Thermo Fisher Scientific) with a uPAC Trapping column (Thermo Scientific) and a 50 cm uPAC Neo HPLC column (Thermo Scientific). For peptide separation and elution, mobile phase A was 0.1% formic acid (FA) in water and mobile phase B was 0.1% FA in acetonitrile. Peptides were injected onto the trap column at 10 µL/min for 3 minutes using the loading pump. Initially the nanoflow rate was set at 0.75 µL/min and 2% mobile phase B while the peptides were loaded onto the trap column, at 2.8 minutes the solvent composition was changed to 10% mobile phase B. At 5 minutes the flow rate was dropped to 0.300 µL/min at 12% mobile phase B. A two-step gradient was used from 12% to 20% mobile phase B for 41.8 minutes followed by 20% to 40% mobile phase for 15.9 minutes. The flow rate was then increased to 0.750 µL/min for column washing using seesaw gradients and re-equilibration. Mass spectrometry analysis was performed on an Orbitrap Eclipse (Thermo Fisher Scientific) operated in data-dependent acquisition mode. The MS1 scans were acquired in Orbitrap at 240k resolution, with a 1 x 10^6^ automated gain control (AGC) target, auto max injection time, and a 375-2000 m/z scan range. MS2 targets were filtered for charge states 2-7, with a dynamic exclusion of 60 seconds, and were accumulated using a 0.7 m/z quadrupole isolation window. MS2 scans were performed in the ion trap at a turbo scan rate following higher energy collision dissociation with a 35% normalized collision energy. MS2 scans used a 1 x 10^4^ AGC target and 35 ms max injection time.

### Protein identification and data filtering for sample comparison

Raw MS data files were processed for protein identification and label-free quantification (LFQ) by MaxQuant (version 2.4.1.0) using the Human SwissProt canonical sequence database (3AUP000005640, downloaded June 2023) and common contaminants (streptavidin, trypsin, albumin and the default MaxQuant contaminants). The following parameters were used: specific tryptic digestion up to four missed cleavages, variable modification search for up to 5 modification per peptide including carbamidomethyl cysteine, protein N-terminal acetylation, and methionine oxidation, default match between run parameters and label-free quantification with minimum ratio count of 1. Only unmodified, oxidized or N-terminal acetylated unique peptides were used for protein quantification. The mass spectrometry proteomics data have been deposited to the ProteomeXchange Consortium via the PRIDE partner repository with the dataset identifier PXD055213 (Reviewer account details: Username; Password: wH0VDRmI46tP). Quantified protein intensity data from MaxQuant were imported and analyzed by Perseus (version 2.0.11). Data were first filtered based on categorical column to remove proteins labeled as “Only identified by site”, “Reverse”, and “Potential contaminant”. Then Intensity values were log2 transformed and replicates were grouped in Categorical annotation rows. Data were further filtered to remove proteins without three valid values in at least one group. Missing values were replaced with 1% limit of detection for the total matrix. MAFG vs control groups were compared using two-sided t-test with FDR of 0.05 and S0 of 0 with the default permutation-based FDR correction for multiple t-tests.

### Proximity Ligation Assay (PLA)

Cells were plated in 8-well Chamber Slide system (Thermo Fisher Scientific Cat. # 154534PK) at a density of 50,000 cells per well. After 16–24 hours, each well was washed twice with PBS and fixed with 100% methanol at −20°C for 20 min. After methanol fixation, cells were washed twice with PBS and PLA was performed with the Duolink In Situ PLA (Sigma Aldrich, Cat. # DUO92008) following the manufacturer’s indications. For the TMA, amplification was performed for 2 hours. Antibodies used were MAFG (Abcam, Cat. # ab154318) and MITF (Santa Cruz, Cat. # sc-56725) at 1:400 for cell lines and at 1:200 for the TMA, with PLA anti-rabbit PLUS (Sigma Aldrich, Cat. # DUO92002) and PLA anti-mouse MINUS (Sigma Aldrich, Cat. # DUO92004) probes. All images were taken at 40x by confocal microscopy and the maxIP projections were analyzed. For each image total foci and nuclear foci per cell were counted.

### Immunoblotting

Cells were washed and scraped in PBS, centrifuged, and the pellet was lysed using RIPA buffer containing protease and phosphatase inhibitor cocktail (Thermo Scientific, Cat. # 78440). 20μg of total protein were subjected to SDS-PAGE and Western blot, performed as previously described^15^. Primary antibodies used were MAFG (Thermo Fisher Scientific, Cat # PA5-90907), AXL (Cell Signaling, Cat.no 8661S), MITF (Cell Signaling, Cat.no 12590S), Flag-M2 (Millipore Sigma, Cat # F1804), HSP90 (Cell Signaling, Cat # 4874), Anti-V5 (Thermo Fisher Scientific, Cat # R96025; RRID: AB_2556564) and β-Actin (Invitrogen, Cat. # AM4302).

### Co-Immunoprecipitation

Pierce protein A/G magnetic beads (Thermo Fisher, Cat. # 88803) were washed with 1% filtered BSA-PBS for 1hr at 4°C with agitation. Cells were washed once with PBS, scraped, and centrifuged to collect cell pellets. Cell pellets were lysed using 200 µl EBC lysis buffer (50 mM Tris-HCL, pH 7.4; 150 mM NaCl; 0.5% IGEPAL, 1:1000 Halt protease and phosphatase inhibitor cocktail (Thermo Fisher, Cat. #78440)), incubated at 4°C for 30 minutes with agitation, and centrifuged at 12,000g for 20 minutes to collect the supernatant. Protein concentration was determined by DC protein assay. Pre-washed beads were incubated with antibodies (Normal Rabbit IgG, Millipore Sigma, Cat. #12-370; Normal mouse IgG, Cell Signaling Technology, Cat. #68860L; MAFG, Abcam, Cat. # ab154318; MITF, Millipore Sigma, Cat. # HPA-003259) for 2 hours at RT with end-over-end rotation. 1mg protein lysate was added to antibody-coated beads and incubated overnight at 4°C with end-over-end rotation. Beads were washed 3 times using lysis buffer and once using ice-cold sterile PBS. Magnetically separated beads were resuspended in 20 µl 1x Laemmli buffer and incubated at 350rpm at RT for 2 minutes to elute protein. Supernatant was then subjected to immunoblotting.

### Immunohistochemistry

Tumor tissues were fixed in 10% buffered formalin overnight and dehydrated in 70% ethanol. Tissues were paraffin-embedded, sectioned, and hematoxylin and eosin stained by IDEXX BioAnalytics (Columbia, MO). The tissue sections were de-paraffinized in xylene and rehydrated through an alcohol series. Antigen retrieval was performed by heating the sections in citrate buffer for 10 minutes followed by blocking endogenous peroxidase activity with 3% hydrogen peroxide. Immunohistochemistry was performed using ImmPRESS HRP goat anti-rabbit kit (Vector Laboratories, Cat. # MP-7451). as per the manufacturer’s instructions and then incubated with DAB peroxidase substrate (Cat. # SK4105). The tissue sections were then counter stained in hematoxylin (Vector Laboratories, Cat. # H-3404). Antibodies against Mafg (Abcam, Cat. # ab154318) and Ki-67 (Cell Signaling, Cat. # 12202S) were used for immunohistochemistry.

### ESC-GEMM models and in vivo experiments

All animal experiments were conducted in accordance with an IACUC protocol approved by the University of South Florida. ES cell targeting and generation of chimeras was performed as described previously^15^. Melanoma development was induced in 3–4-week-old MAFG and GFP control chimeras having similar ESC contribution using 25 mg/mL 4-OH Tamoxifen as described previously^15^. Mice were fed 200 mg/kg Doxycycline (Envigo, Cat. # TD180625) *ad libitum*. Experimental mice were euthanized when IACUC-approved clinical endpoints, typically volume of primary tumors, was reached. NSG mice were obtained from JAX (Stock No: 005557) and bred in-house. 6-week-old male and female NSG mice were randomly divided into groups (at least 5 mice per group). 400,000 WM164 or 200,000 SKMel28 melanoma cells were subcutaneously injected into NSG mice, and tumor growth was measured with calipers every 2-3 days. Experimental mice were euthanized when IACUC-approved clinical endpoints, typically volume of primary tumors, was reached.

### Statistical analysis

Statistical analysis was performed using GraphPad Prism software. Survival data were compared by applying the Gehan-Breslow-Wilcoxon test, and all other data were analyzed with the unpaired two-tailed t-test or ordinary one-way ANOVA. A p-value below 0.05 was considered statistically significant. Data represent the mean ± SD of at least two independent experiments performed at least in triplicate.

## SUPPLEMENTARY FIGURE LEGENDS

**Supplementary Figure S1: MAFG overexpression elicits oncogenic effects in melanocytes, melanoma cells, and in a spontaneous mouse melanoma model.** (**A**) Oncoprint of MAFG copy number gains and mRNA expression in samples from The Cancer Genome Atlas skin cutaneous melanoma (TCGA-SKCM) dataset. (**B**) Western blot validating MAFG overexpression in Hermes1 melanocytes and WM164, SKMel28, A375, and WM793 melanoma cells. Ectopic MAFG is shifted up due to the presence of a Myc-DDK tag. (**C,D**) Proliferation (C) and colony formation (D) assays in WM793 (left) and A375 (right) overexpressing MAFG or a GFP control. (**E**) H&E staining and MAFG immunohistochemistry on tumors from BP^MAFG^ and BP^GFP^ mice at endpoint. (**F**) Quantification of MAFG-positive cells per field in tumors from BP^MAFG^ and BP^GFP^ mice. (**G**) Ki67 immunohistochemistry on tumors from BP^MAFG^ and BP^GFP^ mice at endpoint and quantification of Ki67-positive cells per field. (**H,I**) Kaplan–Meier curves showing the tumor-free survival (H) and overall survival (I) of BCC^GFP^ (n = 8) and BCC^MAFG^ (n = 7) chimeras using the Gehan-Breslow-Wilcoxon test. (**J**) Number of melanomas that developed in the BCC^GFP^ and BCC^MAFG^ chimeras. (**K**) Kaplan–Meier curves comparing the tumor free survival of PP^MAFG^ mice fed a Doxycycline-containing diet (On Dox, n = 10) or a regular diet (Off Dox, n = 4) using the Gehan-Breslow-Wilcoxon test. For in vitro experiments, representative replicates of at least two independent experiments performed in quadruplicate are shown. ns, not significant; * p < 0.05; ** p < 0.01.

**Supplementary Figure S2: The oncogenic effect of MAFG is independent of NRF2.** (**A**) Volcano plots of differentially expressed genes identified by RNA sequencing comparing GFP and MAFG overexpression in WM164 (left), SKMel28 (center) and A375 (right) cells. (**B**) Comparison of the 926 differentially expressed genes in WM164 cells with previously published NRF2 gene expression signatures. (**C**) qRT-PCR showing the expression of *MAFG*, *NRF2*, and a canonical NRF2 target gene (*PRDX1*) in response to NRF2 silencing in WM164 cells overexpressing MAFG or GFP. Cells transfected with non-targeting siRNA were used as control. G, GFP; M, MAFG. (**D,E**) Proliferation (D) and colony formation (E) assays of WM164 cells overexpressing MAFG or GFP following the silencing of NRF2. (**F**) Luciferase assay using a transcriptional reporter (6xARE-Luciferase). WM164 cells overexpressing MAFG or GFP following the silencing of NRF2 are shown on the left and parental WM164 overexpressing a constitutive active NRF2-T80K are shown as a positive control on the right. For in vitro experiments, representative replicates of at least two independent experiments performed in quadruplicate are shown. ns, not significant; * p < 0.05; ** p < 0.01.

**Supplementary Figure S3: MAFG promotes a phenotype switch.** (**A**) Pathway analysis of the differentially expressed genes identified by RNA sequencing comparing GFP and MAFG overexpression in WM164. Turquoise, GeneOntology; Orange, Reactome; Blue, Elsevier; Purple, KEGG. (**B**) qRT-PCR showing *MITF* expression upon MAFG overexpression in WM164 and SKmel28 cells (**C**) UMAP graph showing MAFG expression distribution (left) in human melanoma cell clusters representing different phenotypic states (right) from Pozniak et al., 2024. (**D**) UMAP graph showing MAFG expression distribution (left) in mouse melanoma cell clusters representing different phenotypic states (right) from Karras et al., 2022. (**E**) Western blot showing levels of MITF and AXL in two MITF^hi^ (WM164 and SKMel28) and two MITF^lo^ (A375 and WM793) melanoma cell lines. (**F**) qRT-PCR showing expression of *MITF*, *AXL*, and the MITF target genes *PMEL* and *TYR* in two MITF^hi^ (WM164 and SKMel28) and two MITF^lo^ (A375 and WM793) melanoma cell lines. (**G**) Heatmap of the phenotypic state distribution of two MITF^hi^ (WM164 and SKMel28) and two MITF^lo^ (A375 and WM793) melanoma cell lines obtained by analyzing previously published RNAseq (Vera, Bok et al., 2021) (GSE148552).

**Supplementary Figure S4: MAFG interacts with MITF.** (**A**) Western blot showing the expression of the TurboID-MAFG fusion construct in WM164 cells blotting for MAFG (left) or V5-Tag (right). (**B**) Biotin proximity labeling using N-terminal TurboID-MAFG fusion construct in WM164 cells blotting for Streptavidin-HRP. (**C**) Images of proximity ligation assays in WM164, SKMel28 and A375 cells overexpressing either GFP or MAFG. (**D**) Correlation between the percent of PLA positive nuclei per core and the average number of PLA foci per nucleus in a melanoma metastasis TMA.

**Supplementary Figure S5: MITF is required for the MAFG oncogenic effects.** (**A**) Western blot validating the silencing of MITF in WM164 and SKMel28 cells overexpressing MAFG or GFP. (**B,C**) Proliferation (B) and colony formation (C) assays in SKMel28 cells overexpressing MAFG or GFP following MITF silencing. GFP-siNT vs. MAFG-siNT: * p < 0.01; ** p < 0.001; *** p < 0.0001. GFP-siNT vs GFP-siMITF: &&&, p < 0.001. MAFG-siNT vs MAFG-siMITF: ###, p < 0.001. (**D**) Western blot validating the silencing of MITF with two doxycycline-inducible shRNAs in WM164 cells overexpressing MAFG. (**E,F**) Proliferation (E) and colony formation (F) assays showing the effect of MITF silencing with two doxycycline-inducible shRNAs on WM164 cells overexpressing MAFG. ns, not significant; * p < 0.05; ** p < 0.01; *** p < 0.001. For in vitro experiments, representative replicates of at least two independent experiments performed in quadruplicate are shown.

**Supplementary Figure S6: MAFG impacts MITF target gene binding and transactivation:** (**A**) IGV plots for *SLC7A8* (left) and *SVIL* (right), which harbor overlapping peaks and where MAFG overexpression increases MAFG occupancy without significantly affecting MITF occupancy. (**B**) Expression levels of *SLC7A8* and *SVIL* from the RNAseq shown in Figure 2. (**C**) IGV plot for *MLANA* showing MAFG and MITF overlapping peaks. (**D**) IGV plot for *TYR* showing overlapping MAFG/MITF sites with no differential occupancy, and a unique MAFG site with increased occupancy. ### false discovery rate (FDR) < 0.001.

**Supplementary Figure S7: MAFG impacts MITF target gene binding and transactivation.** (**A**) Western blot showing the expression of the MAFG∼MITF tethered dimer construct in WM164 and SKMel28 cells. (**B**) qRT-PCR showing expression of *AXL* in WM164 and SKMel28 cells overexpressing the tethered MAFG∼MITF dimer. (**C**) Venn diagram showing the overlap of differentially expressed genes upon overexpression of MAFG or the tethered MAFG∼MITF dimer in WM164 cells. (**D**) Pathway analysis of the overlapping genes in (C). (**E**) Expression changes of the melanoma phenotype-associated gene signatures in WM164 cells expressing the tethered MAFG∼MITF dimer or GFP. For in vitro experiments, representative replicates of at least two independent experiments performed in quadruplicate are shown. ** p < 0.01; *** p < 0.001; **** p < 0.0001.

**Supplementary Figure S8: MAFF and MAFK possess oncogenic potential in melanoma.** (**A**) Western blot showing the overexpression of Flag-MAFG, Flag-MAFK and Flag-MAFF and their effect on AXL and MITF levels in WM164 cells. (**B,C**) Proliferation (B) and colony formation (C) assays of WM164 cells overexpressing MAFG, MAFF, MAFK, or GFP. (**D**) Co-immunoprecipitation of MITF in WM164 cells overexpressing MAFG, MAFF, or MAFK. Quantification of the relative binding of MAFG, MAFF, and MAFK to MITF is shown on the right. For in vitro experiments, representative replicates of at least two independent experiments performed in quadruplicate are shown. ** p < 0.01; *** p < 0.001; **** p < 0.0001.

## AUTHOR CONTRIBUTION

O.V. and F.A.K. conceived the project and O.V., M.M., Z.S.V., and F.A.K. designed experiments. O.V., M.M., and Z.S.V. performed experiments with help from K.W., X.X., S.R., N.M., M.C., B.P., A.A., I.B., Q.L., H.M., and Y.K. J.L.M provided and scored tissue microarrays. K.Y.T performed histopathological evaluations. I.B. and M.B.M performed biotin proximity labeling analysis. Q.L. and E.K.L assisted with proximity ligation assays. X.Y. performed bioinformatic and statistical analyses. M.B.M, E.K.L, X.Y., I.I., and F.A.K. supervised the studies and acquired funding.

## ACKNOWLEDGEMENTS

We thank G. DeNicola for plasmids and K. Smalley, G. DeNicola, and members of the Karreth lab for helpful discussions. O.V.P. and IIC were supported by Instituto de Salud Carlos III and co-funded by the European Union under Grants PI18/00050, PI21/00145, and CD22/00040. X.X. received support from a Miles for Moffitt Postdoc Award and a Melanoma Research Foundation Career Development Award (1068914). I.B. was supported by a T32 training grant (CA113275). M.B.M was supported by NIH grant R01CA244236. F.A.K. received funding from the American Cancer Society (RSG-21-087-01), Melanoma Research Alliance (https://doi.org/10.48050/pc.gr.75702), and NIH grants R01CA259046 and R21CA256141. This work was also supported by the Gene Targeting Core, Bioinformatics and Biostatistics Shared Resource, Molecular Genomics Core, Analytical Microscopy Core, which are funded in part by Moffitt’s Cancer Center Support Grant (P30CA076292).

